# Age-related differences in ocular features of a naturalistic free-ranging population of rhesus macaques

**DOI:** 10.1101/2022.07.29.501993

**Authors:** Arthur G Fernandes, Palaiologos Alexopoulos, Armando Burgos-Rodriguez, Melween I Martinez, Cayo Biobank Research Unit, Mark Ghassibi, Ilya Leskov, Lauren J N Brent, Noah Snyder-Mackler, John Danias, Gadi Wollstein, James P Higham, Amanda D Melin

## Abstract

**Purpose:** Rhesus macaques (*Macaca mulatta*) are the premier nonhuman primate model for studying human health and disease. We aimed to investigate if age was associated with ocular features of clinical relevance in a large cohort of free-ranging rhesus macaques from Cayo Santiago, Puerto Rico.

**Methods:** We evaluated 120 rhesus macaques (73 males, 47 females) aged from 0 to 29 years old (mean±SD: 12.6±6.4) from September to December, 2021. The ophthalmic evaluation included IOP assessment, corneal pachymetry, anterior segment biomicroscopy, A-Scan biometry, automated refraction, and fundus photography after pupil dilation. The effects of age on the outcome variables were investigated through multilevel mixed-effects models adjusted for sex and weight.

**Results:** On average, IOP, pachymetry, axial length, and automated refraction spherical equivalent were 15.47±2.47 mmHg, 474.43±32.21 μm, 19.49±1.24 mm, and 0.30±1.70 D, respectively. Age was significantly associated with pachymetry (Coef.= -1.20; 95%CI: -2.27 to -0.14; p=0.026), axial length (Coef.= 0.03; 95%CI: 0.01 to 0.05; p=0.002), and spherical equivalent (Coef.= -0.12; 95%CI: -0.22 to -0.02; p=0.015). No association was detected between age and IOP. The prevalence of cataracts in either eye was 10.83% (95% CI: 6.34 – 17.89%) and was significantly associated with age (OR= 1.20; 95%CI: 1.06 – 1.36; p=0.004). Retinal drusen in either eye was observed in 15.00% (95% CI: 9.60 – 22.68%) of the animals, which was also significantly associated with age (OR=1.14; 95%CI: 1.02 – 1.27; p=0.020).

**Conclusions:** Rhesus macaques exhibit age-related ocular associations similar to those observed in human aging, including decreased corneal thickness, increased axial length, myopic shift, and higher occurrence of cataract and retinal drusen.

## INTRODUCTION

Over 4% of the wordwide population is estimated to live with moderate to severe visual impairment or blindness. ^1^ Among the four main causes, cataract, glaucoma, and age-related macular degeneration (ARMD) are conditions strongly associated with aging. ^2^ Blindness due to cataract is a reversible condition and the underlying mechanisms associated with the disease development and progression are mostly known. ^3^ Conversely, the environmental and genetic contributions to diseases such as glaucoma and ARMD are not fully understood, in large part due to the lack of high-quality, medically-relevant, unbiased data, as well as confounders introduced by medical interventions as well as social and environmental variation among subjects. ^4-6^ Given these limitations, the evaluation of animal models in consistent environments is a valuable alternative to overcome the gaps and promote better understanding of these diseases. Studies of primates are especially valuable for their translational potential to humans due to shared general biology and physiology, including many shared ocular features. ^7^

Rhesus macaques (*Macaca mulatta*) are the premier nonhuman primate (NHP) model for studying human health and disease. ^8^ Members of the genus Macaca have over 90% DNA sequence similarity and highly conserved protein sequences with humans. ^8^ The rhesus macaque lifespan is approximately 3–4 times shorter than that of humans, which provides an advantage for longitudinal research, as age-related changes occur over a ca. four-fold shorter time span. ^9^ Importantly, the aging process of rhesus monkeys is highly similar to that of humans, including declines in physical health, physiological integrity, brain function, and peripheral immune regulation. ^9-11^

Ocular studies of rhesus macaques have shown close anatomical association between NHP and human visual systems, including features of the optic nerve head, retina, and lens. ^8,12,13^ While those similarities are well established, detailed data on how aging impacts ocular structures in macaques remain limited to a few studies based on small sample sizes of captive animals. Data from a free-ranging, naturalistic population, free from medical interventions, hold considerable promise to inform our understanding of the similarities and differences between the NHP and human visual system and enhance translational outcomes. The population of rhesus macaques residing on the island of Cayo Santiago, a 15.2-hectare island located 1 km from the eastern coast of Puerto Rico, have provided a unique resource for medical and social research for over 80 years. ^14^ The macaques live under naturalistic circumstances and are free to socialize with others in the population. ^9^

The purpose of the current study is to quantify age-related differences in ocular features in a large cohort of free-ranging rhesus macaques from Cayo Santiago. We specifically aim to understand the association of age with ocular structure, as well as with other age-related features commonly seen in humans as cataracts and retinal drusen.

## METHODS

### Study population

The Cayo Santiago population of rhesus macaques was established in the late 1930s and started with 409 founders imported from India. Since then, the macaques have flourished, with the population now approaching 2000 individuals. The population is managed by the Caribbean Primate Research Center (CPRC) at the University of Puerto Rico. The CPRC maintains a detailed census of the population, recording the date of birth of each monkey, and provisions them daily with monkey chow and water. Aside from tetanus inoculation and tattoos that are administered when animals are approximately one year old, no medical interventions are administered to the individuals. The animals at Cayo Santiago are free to aggregate into social units, supplement their diets with insects and leaves found on the island, are exposed to natural weather conditions, and interact in affiliative, agonistic, and reproductive behaviors with other members of the population. ^9^ Each individual is recognized and tracked by tattoos, ear notches, and facial features.

We performed comprehensive eye exams on 120 rhesus macaques (73 male and 47 female) from September to December, 2021. To consider biological variables that may impact ocular features, we collected data on age and weight of all study individuals (Table 1). Males and females in our dataset did not differ significantly in age (z=0.680; p=0.4964)), but males were significantly heavier than females (z=-7.371; p<0.0001).

**Table 1.** Rhesus macaque age and weight (mean with standard deviation; SD) by sex

| **Variables** | **<u>Males</u>** | **<u>Females</u>** | **<u>All</u>** |
| --- | --- | --- | --- |
|  | *mean±SD* | *mean±SD* | *mean±SD* |
| **Age (years)** | 12.40±5.60 | 12.80±7.62 | 12.56±6.44 |
| **Weight (lbs)** | 21.70±7.07 | 13.17±6.02 | 18.36±7.86 |

### Eye examination procedures

All the procedures complied with the guidelines of the Association for Research of Vision and Ophthalmology. Animals were anesthetized using Ketamine HCL (100mg/ml) and Xylazine (100mg/ml) following the protocol approved by the animal welfare committee of the University of Puerto Rico (IACUC #A400117) and of the University of Calgary (AC19-0091).

The ophthalmic evaluation included IOP, corneal pachymetry, anterior segment biomicroscopy, and A-Scan biometry. Automated refraction, and fundus photography were performed after pupil dilation. All in-vivo evaluation and testing were conducted by one experienced ophthalmic technologist (AF) with experience in NHP handling. We measured IOP using a Tonovet tonometer (iCare, Vantaa, Finland). Central corneal thickness was measured using Pachmate 2 ultrasonic pachymeter (DGH Technologies, Exton, PA). Anterior segment biomicroscopy was performed using a handheld slit lamp (MicroClear, Jiangsu, China) with focus on cataract diagnosis, considering any type of lens opacification. The A-Scan was performed using VuPad (Sonomed Escalon, Lake Success, NY, USA) and provided measurements of axial length, anterior chamber depth, lens, and vitreous chamber. Pupils were dilated by using a drop of tropicamide 1% and one drop of cyclopentolate 1%. Static refraction was performed with the automated refractor KR 7000S (Topcon, Tokyo, Japan) and provided both refraction and keratometry information. Finally, fundus photography was performed using the posterior segment module from the TRC-NW400 camera (Topcon, Tokyo, Japan) and images were later analyzed by retina specialists (MG and IL).

Both pachymetry and A-Scan were performed after the instillation of one anesthetic drop tetracaine 0.5% in each eye to guarantee the animal comfort. A-Scan was not performed in individuals younger than 3 years old due to the equipment limitation.

Automated refraction was performed at least 20 minutes from the administration of pupil dilation drops in individuals older than 7 years old. The spherical equivalent was calculated as the refraction spherical component summed to half of the cylinder component. A rigid gas permeable contact lens (Boston EO, Boston, MA) was fitted to each eye to improve the fundus photography image quality. Each image was analyzed by two independent retina experts to determine retinal drusen presence. After the exams the animals were kept monitored until the next day and were evaluated by the veterinarian before being released.

### Statistical methods

We used Stata/SE Statistical Software, Release 14.0, 2015 (Stata Corp, College Station, TX, USA) for statistical analyses. Frequency tables were used for descriptive analysis. We evaluated the effects of age on IOP, pachymetry, axial length, and auto-refraction spherical equivalent values through multilevel mixed-effects models adjusted for sex and weight. We investigated the effects of age on cataract and retinal drusen occurrences by multiple logistic regression adjusted for the sex and weight. P values ≤0.05 were considered statistically significant.

## RESULTS

The mean IOP, corneal thickness, axial length, and automated refraction spherical equivalent are presented in Table 2. We found significant effects of age on corneal thickness, axial length, and spherical equivalent (Figure 1). For each year of age, the corneal thickness decreased by 1.20 μm, the axial length increased 0.03 mm, and the spherical equivalent decreased 0.12 diopters. There was also a main effect of sex on IOP and axial length where males had mean IOP that was 0.98mmHg lower than females, and axial lengths that were 0.43 mm longer than females. Moreover, there was a significant effect of weight on the intraocular pressure so that for each 1 increasing pound of weight, the intraocular pressure increased by 0.21 mmHg (Table 3).

**Table 2.** Features of ocular structures measured in a free-ranging population of rhesus macaques

| Variable | Mean±std | Range |
| --- | --- | --- |
| IOP (mmHg) | 15.47±2.47 | 9 – 23 |
| Pachymetry (μm) | 473.06±32.16 | 394 – 551 |
| **A-Scan (mm)** |  |  |
| ACD | 3.45±0.24 | 2.59 – 4.06 |
| Lens | 3.31±0.25 | 2.81 – 4.05 |
| Vitreous | 12.71±0.54 | 11.47 – 14.11 |
| AL | 19.46±0.56 | 18.11 – 20.81 |
| **Refraction (D)** |  |  |
| Spheric | 1.19±1.77 | -6.75 – 6.00 |
| Cylinder | -1.79±1.53 | -6.75 – -0.25 |
| Spherical Equivalent | 0.30±1.70 | -8.38 – 3.50 |
| K1 | 51.21±2.76 | 42.50 – 55.87 |
| K2 | 53.21±2.46 | 48.37 – 60.37 |
| KM | 52.41±2.74 | 47.12 – 63.75 |

*ACD: Anterior Chamber Depth; AL: Axial Length*

**Figure 1.**
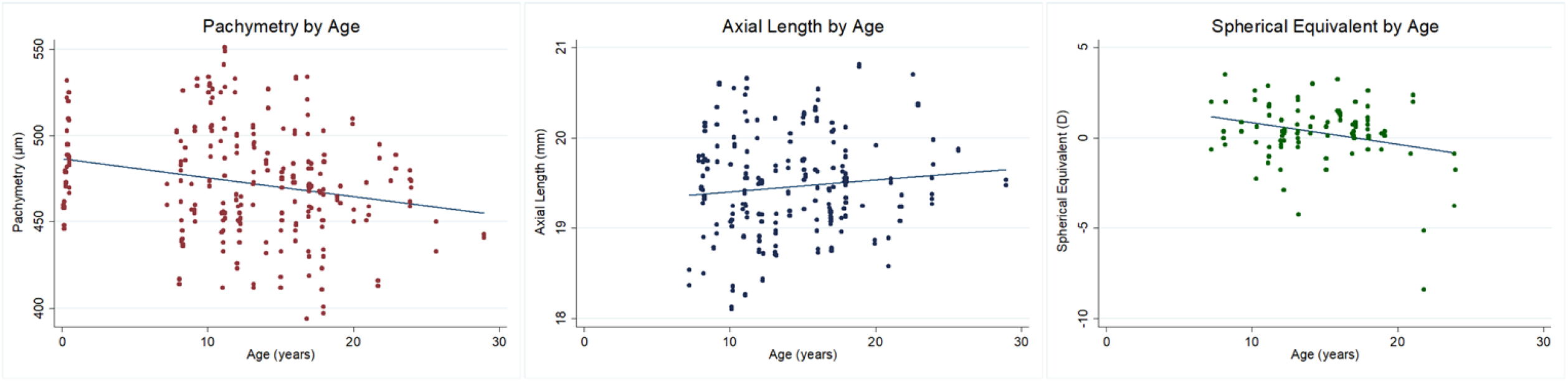
Association of age and: A) Pachymetry; B) Axial length; and C) Spherical equivalent.

**Table 3.** Multilevel mixed-effects model analysis for IOP, pachymetry, axial length, and spherical equivalent

|  | IOP |  | PACHYMETRY |  | AXIAL LENGTH |  | SPHERICAL EQUIVALENT |  |
| --- | --- | --- | --- | --- | --- | --- | --- | --- |
| *Variable* | *Coefficient (95% CI)* | *p-value* | *Coefficient (95% CI)* | *p-value* | *Coefficient (95% CI)* | *p-value* | *Coefficient (95% CI)* | *p-value* |
| **Sex** |  |  |  |  |  |  |  |  |
| Female | Ref. | --- | Ref. | --- | Ref. | --- | Ref. | --- |
| Male | -0.98 (-1.94 to -0.02) | **0.046** | 11.21 (-3.02 to 25.44) | 0.123 | 0.43 (0.11 to 0.76) | **0.008** | 0.46 (-1.03 to 1.95) | 0.549 |
| **Age** | -0.05 (-0.12 to 0.02) | 0.156 | -1.20 (-2.27 to -0.14) | **0.026** | 0.03 (0.01 to 0.05) | **0.002** | -0.12 (-0.22 to -0.02) | **0.015** |
| **Weight** | 0.21 (0.14 to 0.28) | **<0.001** | -0.24 (-0.79 to 1.26) | 0.647 | 0.02 (-0.01 to 0.06) | 0.219 | -0.02 (-0.18 to 0.14) | 0.785 |

Cataract (Fig. 2A) in either eye was observed in 13 animals (7 unilateral and 6 bilateral cases) representing a frequency of 10.83% (95% CI: 6.34 – 17.89%). Retinal drusen (Fig. 2B) in either eye was observed in 18 animals (10 unilateral and 8 bilateral cases) representing a frequency of 15.00% (95% CI: 9.60 – 22.68%).

**Figure 2.**
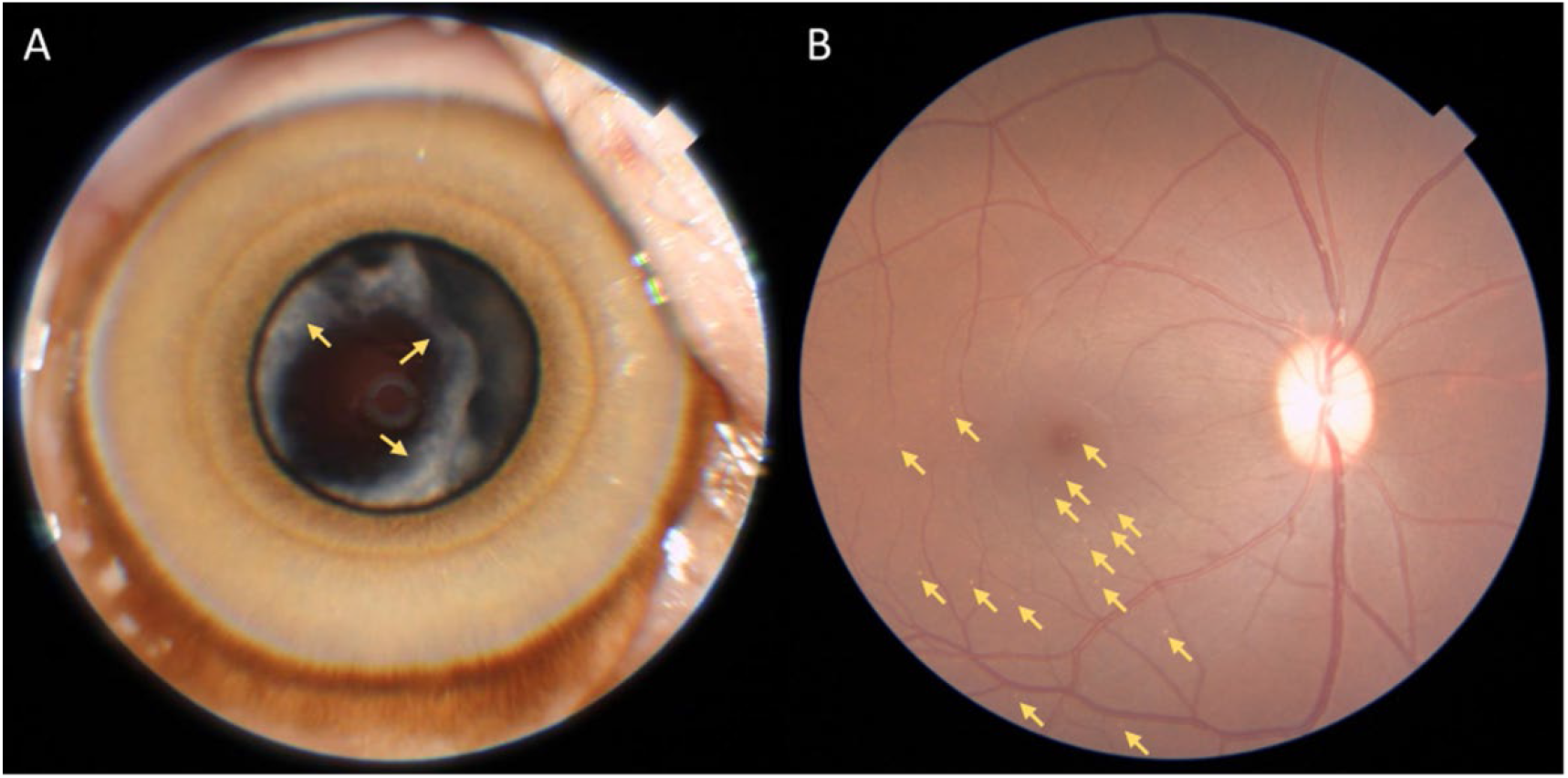
A) Cataract, right eye, female, 16.4 years old. Yellow arrow indicates cortical lens opacities. B) Retinal drusen, right eye, male, 19.1 years old. Yellow arrows indicate drusen concentrated on macular and temporal inferior regions.

Older individuals were more likely to have cataract (OR= 1.20; 95%CI: 1.06 – 1.36; p=0.004) and retinal drusen (OR=1.14; 95%CI: 1.02 – 1.27; p=0.020) (Table 4). The mean number of drusen on the 26 affected eyes was 8.9±7.6. Despite the association between the presence of drusen and age, no significant associations between number of drusen and sex, age, or weight were observed (p>0.05).

**Table 4.** Multiple logistic regression for cataract and retinal drusen in either eye

|  | **CATARACT** |  | **RETINAL DRUSEN** |  |
| --- | --- | --- | --- | --- |
| *Variable* | *Odds Ratio (95% CI)* | *p-value* | *Odds Ratio (95% CI)* | *p-value* |
| **Sex** |  |  |  |  |
| Female | Ref. | --- | Ref. | --- |
| Male | 0.45 (0.07 – 2.77) | 0.389 | 1.54 (0.30 – 7.94) | 0.605 |
| **Age** | 1.20 (1.06 – 1.36) | **0.004** | 1.14 (1.02 – 1.27) | **0.020** |
| **Weight** | 0.99 (0.83 – 1.20) | 0.957 | 1.05 (0.91 – 1.22) | 0.505 |

Ocular abnormalities that were not age-related were less frequent and included corneal scars/leucoma (2.50%; e.g Figure 3), eyelid laceration (0.83%), conjunctiva (0.83%), and iris (0.83%) nevus; the first two likely due to trauma events and/or infections that occurred during typical social behaviors.

**Figure 3.**
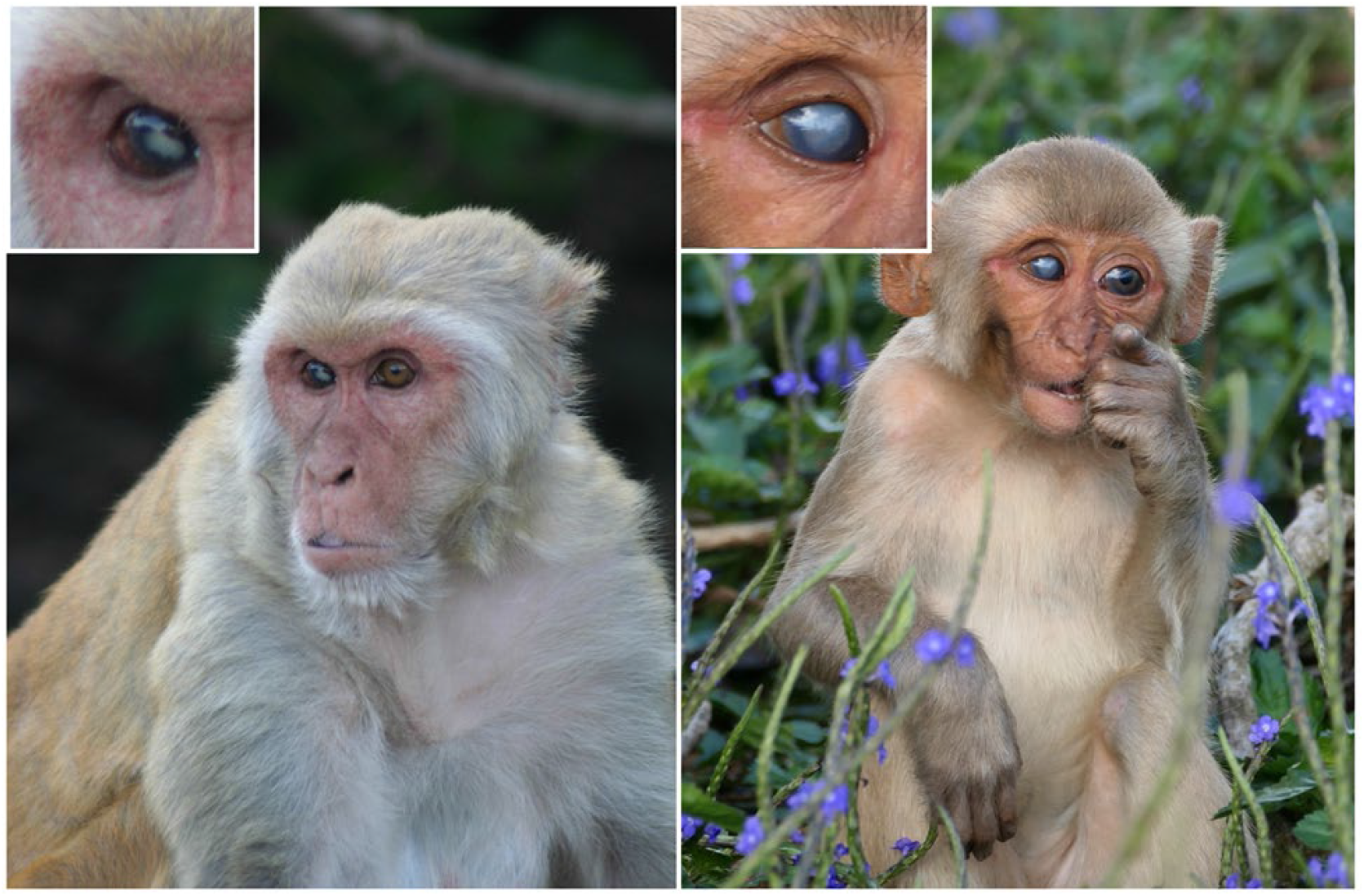
An adult (A) and a juvenile (B) rhesus macaque (*Macaca mulatta*) on Cayo Santiago with visible leucoma in the right eyes. The sources of the leucomas are unknown but they were likely to be caused by trauma from their environment or conspecifics.

## DISCUSSION

We investigated age-related ocular features in a large cohort of free-ranging rhesus macaques. We found that older individuals had thinner corneas, longer axial length, myopia, and higher incidence of cataract and retinal drusen. Below we discuss and contextualize these findings in more detail.

Rhesus macaques age in a manner that closely resembles humans, including similar periods of development, maturation, reproduction and senescence, albeit within a four-fold shorter time span. ^9^ These monkeys display typical aging hallmarks, such as thinning and greying hair, atrophying skin, declining motor activity, decreasing stature, redistribution of body fat, and also exhibit pathological signs of age-related diseases such as cancer, cardiovascular disease, and metabolism disorders. ^9,10,15-17^ With continual increases in human longevity, conditions such as glaucoma and age-related macular degeneration are expected to be major causes of vision impairment and blindness, ^18,19^ highlighting the importance of identifying a good animal biomedical model for these and related ocular conditions.

Our study shows age-related differences in corneal thickness, axial length, and refraction in rhesus macaque eyes. While these features are not necessarily associated with disease, per se, they contribute to a growing understanding of changes that happen in the eye as we age. Our results indicate a corneal thickness decrease rate of 1.20 μm per year. Human studies also show decline in corneal thickness, but extensive variation in the estimated rates is present, due to differing methodologies and devices used for measurement, with rates ranging from 0.30 μm/year when using optical devices ^20-22^ to 1.20 μm/year when using ultrasound-based devices. ^23,24^ Corneal thickening has been attributed to loss of both epithelial and endothelial cells, which has been demonstrated to be associated with the aging process. However, other underlying causes such as changing in overall cell distribution and limbal epithelial stem cells dysfunction are under study. ^20,21,24,25^ Importantly, corneal thickening rate has been found to be significantly greater in glaucoma cases, ^24^ reinforcing the importance of a model with similar corneal characteristics when studying the disease. Regarding axial length, althought the Sorbys’s classical statement indicates that the axial eye growth in humans is completed at the age of 13 years old, recent studies have demonstrated that the eye of those with persistent emmetropia and hyperopia continues to grow between 18 and 22 years, at a slow rate of approximately 0.03–0.04 mm/year. ^26,27^ In our sample of rhesus macaques, we found an axial length increasing rate of 0.03 mm/year, slightly lower than a previous publication in rhesus macaques that reported a rate of 0.05 mm/year, ^28^ a difference possibly due to a broader range of ages in their analysis. Finally, as a consequence of the axial length increase, a myopic shift on the static spherical equivalent is expected to be observed. ^29^ Longitudinal studies with human populations have demonstrated that a 0.1 mm change in axial length correspond to -0.24 D change in myopia. ^30^ Our results, indeed, showed a change rate of -0.12 D/per year, which can also be influenced by cataract formation in older adults.

We found that IOP was not significantly associated with age. This is perhaps surprising given that a recent study on rhesus macaques reported higher IOP values in older individuals. ^28^ However, a previous study on rhesus macaques from Cayo Santiago reports results similar to those we find here, i.e. that age was not a significant predictor of IOP. ^31^ Results from human studies are also variable, with some reports indicating higher IOP, ^32^ lowerIOP, ^33,34^ or no IOP association ^35,36^ in older individuals relative to younger individuals. Finally, we found a positive association between IOP and weight, which is in agreement with expectations and previous research. ^32,37^

In accordance with previous studies in humans, we found cataracts to be strongly associated with aging. ^2^ Several population-based studies in humans have reported increased risk of cataracts ranging from 6% to 24% for each increasing year in age, similar to our finding of 20%. ^2,36,38,39^ Similarly, a higher frequency of retinal drusen is expected in older individuals. ^40,41^ Our findings indicate a frequency of retinal drusen of 15% in the population and its significant association with age, with the odds of having drusen increasing by 9% for each year of age. Previous studies evaluating drusen in rhesus macaques have shown variable results with frequencies ranging from 5 to 61% of the population, ^42-46^ variability potentially explained by several factors such as age and sex distribution, genetic differences among colonies, environment, methods of examination, and criteria used to define drusen occurrence. Retinal drusens may not cause visual impairment per se but are considered one of the main risk factors for age-related macular degeneration, and therefore a target for investigations of the disease process. ^40,41,47^

Biomedical research in ophthalmology is often reliant on rodents and rabbits, due to the advantages of short lifespan, well-known genetic features, low-cost maintenance, and ease handling. ^48,49^ Limitations of such models, however, include differential optic nerve head structure and lamina cribosa and dissimilarities on retinal ganglion cell subtypes when compared to primates, ^28^ which restricts the results translation for human diseases as glaucoma. Moreover, non-primate mammals do not have a macula, ^50,51^ again limiting this model’s potential when considering age-related macular degeneration research. We join other researchers in advocating for the rhesus macaques as a model for eye diseases. Here, we present the most detailed and extensive data to date onIOP, corneal thickness, axial length, refraction, and cataract and retinal drusen occurrence in rhesus monkeys. Importantly, this is the first study of ocular features in a naturalistic free-ranging population. Given that aging and disease development often differ between captive and free-ranging animals, ^9,16,52^ our results might represent the most realistic scenario available, well suited for studying natural history of diseases. Limitations of this study include the fact that axial length and static refraction assessments were not performed in individuals younger than 3 and 7 years old, respectively, which could have underestimated the rates of eye growth and myopia shift per year in the general population; we were not able to perform examinations using optical coherence tomography due to logistic limitations of having the equipment located in situ in the field; and finally, our analysis is relied on structural rather than functional features.

In conclusion, rhesus macaques have age-related ocular changes similar to those observed in human aging, including higher occurrence of cataract and drusen, decreasing corneal thickness, increasing axial length, and myopic shift. These results are further evidence that rhesus macaques are a valuable model for study of age-related eye diseases.

## ACKNOWLEDGMENTS

We thank our colleagues for their important support in data collection on Cayo Santiago site, especially Allysa Arre, Nahiri Barreto, Josué Negrón, Daniel Phillips, and the Caribbean Primate Research Center staff. Support for this research was provided by New Frontiers in Research Foundation (NFRFE-2018-02159), Natural Sciences and Engineering Research Council (RGPIN-2017-03782; ADM), Canada Research Chairs Program (950-231257), National Aging Institute (1R56AG071023, R01AG060931), BrightFocus Foundation (G2020047), and University of Calgary Eyes High Fellowship (AGF).

